# Pachytene piRNAs define a conserved program of meiotic gene regulation

**DOI:** 10.1101/2025.11.05.686807

**Authors:** Zuzana Loubalova, Franziska Ahrend, Daniel Stoyko, Rachel Cosby, Sherry Ralls, Gunter Meister, Todd Macfarlan, Astrid D. Haase

## Abstract

PIWI-interacting RNAs (piRNAs) safeguard genome integrity and fertility across animals and in humans^1-3^. In mammals, pre-pachytene piRNAs execute the ancestral program of transposon silencing, whereas a second, mammalian-specific class functions during meiosis^4^. These pachytene piRNAs are indispensable for spermatogenesis^3^, yet their sequence diversity, rapid evolution, and lack of obvious complementarity have long obscured their primary targets and mode of action^5^. Here, we show that groups of pachytene piRNAs converge on single mRNAs with near-perfect complementarity, evoking the specificity and efficacy of classical RNA interference^6^. These mRNA-targeting piRNAs originate from pseudogene fragments within discrete piRNA clusters, establishing an unexpected one-to-one relationship: a given cluster encodes thousands of piRNAs that collectively silence a single gene. Deleting such piRNA clusters or their embedded pseudogene fragments, eliminates the gene-targeting piRNAs, derepresses the target mRNA, and causes spermatogenic defects. Comparative genomics reveals a simple, unifying logic, whereby rapidly evolving cluster–target pairs coexist with a conserved, primate-specific regulon maintained over tens of millions of years. These findings provide a mechanistic framework for pachytene piRNA action, repositioning them as precise gene regulators during meiosis and highlighting a uniquely preserved piRNA–target pair in primates.

---

Pachytene piRNAs are indispensable for spermatogenesis, yet their molecular functions remain largely elusive^4,7^. They originate from hundreds of piRNA precursor genes within discrete genomic loci known as piRNA clusters^8-10^. Millions of piRNAs are parsed from long single-stranded precursors by the endonuclease PLD6 (MitoPLD) and are loaded onto PIWIL1^9-12^. In mice, loss of PIWIL1 (MIWI) abolishes the pachytene piRNA pathway and causes male sterility with post-meiotic arrest at the round spermatid stage^3^. MIWI’s function depends on its RNase H–like slicer activity, characteristic of Argonaute proteins, suggesting a role in cleaving piRNA-complementary target-transcripts^13^. However, direct functional targets have remained unknown. The extreme sequence diversity of pachytene piRNAs, their lack of obvious complementarity to annotated transcripts, and the absence of defined pairing rules have hindered target prediction, leaving the molecular mechanisms by which pachytene piRNAs act unresolved^14-17^.

Here, we developed a target-centered strategy to examine how cellular mRNAs engage with the collective piRNA pool, rather than searching for targets of individual piRNAs. While most pachytene piRNAs lack substantial complementarity to other transcripts, we found that approximately 1.6% match mature mRNAs with ≤1 mismatch across nt 2-21 (**Fig. 1a and Extended Data Fig. 1a**). Although this represents only a small fraction of the total piRNA population, it corresponds to an enormous absolute number, roughly 160,000 molecules per cell, based on the exceptional abundance of piRNAs in germ cells^17,18^. Notably, more than 12% of these gene-targeting piRNAs (∼19,000 molecules per cell) converge on a single mRNA, the histone code reader *Spindlin1* (*Spin1*)^19^, predicting hundreds of potential cleavage sites across more than one kilobase (kb) of its coding sequence (CDS) and 3′ untranslated region (3’UTR) (**Fig. 1a insert, b**). The density and precision of these matches suggest potential for direct piRNA-guided target-slicing by MIWI^20^.

**Figure 1.**
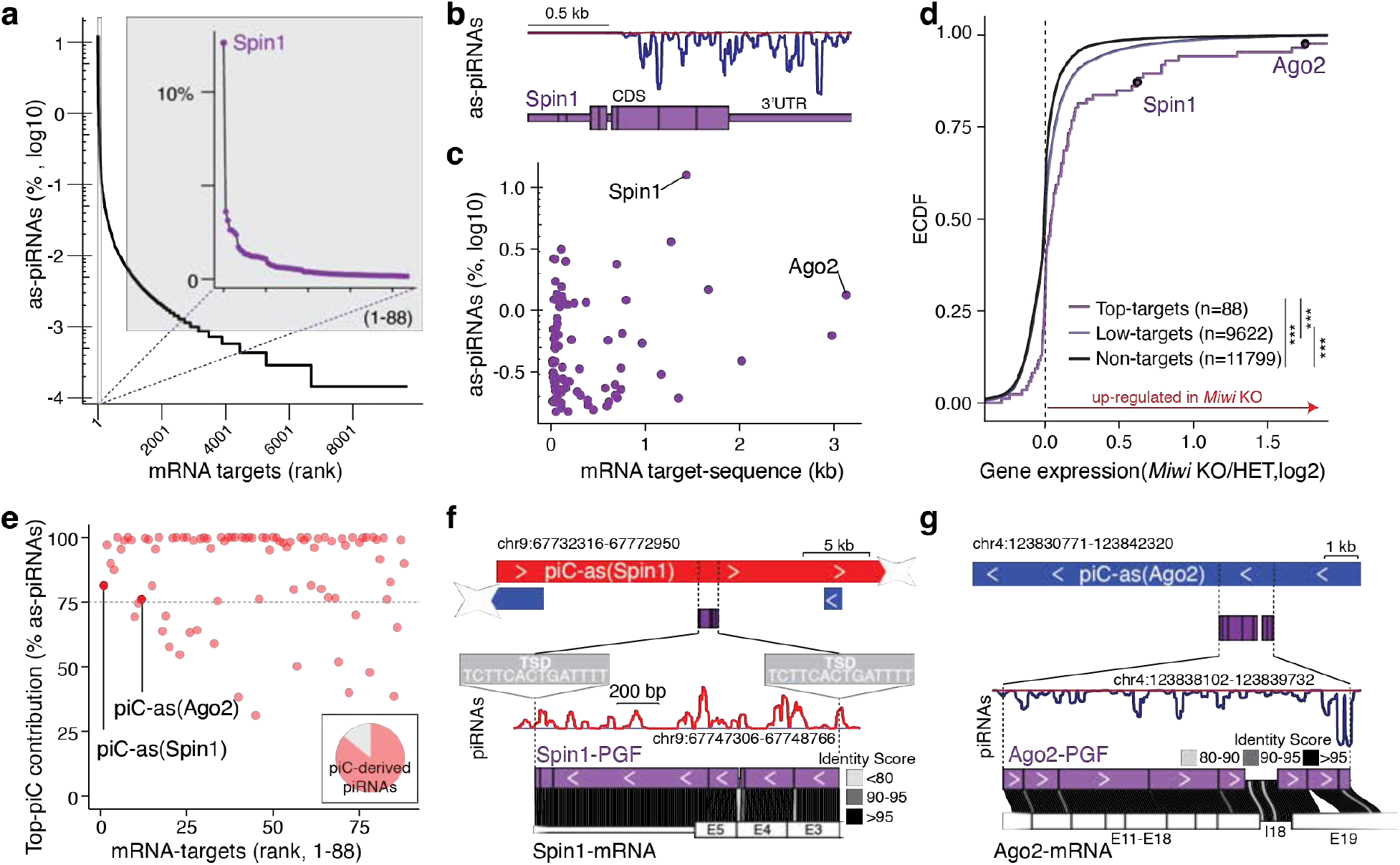
Pachytene piRNAs from discrete clusters converge on single mRNAs across extended CDS and 3′ UTR. **a**, mRNA targets ranked by the fraction of antisense (as-)piRNAs among all mRNA-targeting piRNAs. **Inset:** the top 88 targets account for 70% of all mRNA-targeting piRNAs; the top gene, *Spin1*, receives 12%. **b**, Antisense piRNA coverage across *Spin1* exons. **c**, Relationship between mRNA length (kb) and the fraction of converging as-piRNAs (log_10_ of % of all mRNA-targeting piRNAs) for the top targets (70^th^ percentile; n=88). *Spin1* and *Ago2*, the most strongly and the most extensively targeted genes, respectively, are indicated. **d**, Empirical cumulative distribution function (ECDF) of expression changes (log_2_ fold change) in late spermatocytes from *Miwi* knockout (KO) versus heterozygous (HET) mice (RNA-seq SRR610673–8). Groups: top targets (70^th^ percentile; n = 88), lower-ranked targets (ranks > 88; n = 9,622), and non-targets (n = 11,799). Top targets, including *Spin1* and *Ago2*, skew toward up-regulation in *Miwi* KO (Kolmogorov–Smirnov test, \*\*\**P* < 0.001). **e**, Contribution of as-piRNAs from the dominant (top) mRNA-targeting piRNA cluster (piC) for each target (top 88). Most piRNAs targeting *Spin1* or *Ago2* originate from a single cluster, *piC-as(Spin1)* or *piC-as(Ago2)*, respectively. **Inset:** 86% of mRNA-targeting piRNAs originate from piRNA clusters. **f**, *piC-as(Spin1)* contains a processed *Spin1* pseudogene fragment (*Spin1-*PGF) flanked by target-site duplications (minus strand). piRNA coverage is antisense to the PGF (plus strand, red). The *Spin1*-PGF comprises exons 3–5 with >90% identity to the cognate mRNA. **g**, *piC-as(Ago2)* highlighting the partly processed *Ago2*-PGF with antisense piRNA coverage (minus strand, blue). The fragment aligns to exons 11–19 and intron 18 of *Ago2* with high sequence identity.

Similar to *Spin1*, most mRNA-targets were engaged by hundreds of piRNA sequences across extensive regions spanning their CDS and 3’ UTR (**Fig. 1c, Extended Data Fig. 1a**). *Argonaute-2* (*Ago2*), the major mammalian microRNA (miRNA) partner, showed the longest target space (**Fig. 1c, Extended Data Fig. 1b**). Although piRNA-targeting rules remain debated, with some studies suggesting a need for extensive and others minimal homology, our observation that hundreds of piRNAs converge on targeting individual mRNAs with near-perfect complementarity leaves little doubt that these targets could be efficiently regulated^15,20,21^.

To probe whether piRNAs regulate their predicted targets, we compared the expression of top-ranked (strongly targeted) and lower ranked (weakly targeted) mRNAs, as well as non-targets, in *Miwi* knockout (KO) spermatocytes^22^. Transcripts with the strongest predicted piRNA targeting displayed significantly higher mRNA levels than low-ranking targets or non-targets in *Miwi* KO spermatocytes relative to heterozygous controls (**Fig. 1d**). Among these, *Spin1* was modestly upregulated (<2-fold), whereas *Ago2* showed pronounced increase in mRNA levels (>3-fold) (**Fig. 1d**). However, as expected, direct piRNA targets were not necessarily among the most upregulated genes, since widespread secondary effects accompany the severe spermatogenic defects in *Miwi* KO animals (**Fig. 1d**).

Next, we traced the origin of these gene-targeting piRNAs to determine whether they arose from antisense transcripts of their corresponding genes (cis-regulation) or from distant loci acting in trans. Most piRNAs arose from genomic intervals distant from their target genes, consistent with regulation in trans. Despite their extensive complementarity to target mRNAs, their perfect genomic matches mapped elsewhere. Notably, these mRNA-targeting piRNAs were not scattered across the genome but originated from distinct regions within annotated piRNA clusters (piC)^23^, with a single cluster typically producing the majority of piRNAs targeting a given gene (top-piC) (**Fig. 1e-g, Extended Data Fig. 1d**). Within clusters, the mRNA-targeting piRNAs originated from pseudogene fragments (PGFs) with high sequence identity to their parental genes, providing extensive antisense information for target recognition.

A striking example is the *Spin1*-targeting piRNA population, of which 81% originate from a single piRNA cluster on chromosome 9, designated *piC-as(Spin1)* (**Fig. 1e**). *piC-as(Spin1)* produces antisense *Spin1* piRNAs from a processed pseudogene insertion that selectively targets *Spin1* exons 3-5 (**Extended Data Fig. 1e**). The *Spin1* pseudogene fragment (PGF) is 5’ truncated and flanked by a target-site duplication, features characteristic of a LINE–mediated retrotransposition event^24^. Sequence identity analysis between the PGF and the cognate *Spin1* mRNA revealed high similarity, consistent with extensive targeting potential (**Fig. 1f**). Similarly, 76% of *Ago2*-targeting piRNAs originate from *piC-as(Ago2)* on chromosome 4 (**Fig. 1e, g, Extended Data Fig. 1f**). Together, these findings establish a one-to-one regulatory relationship between individual piRNA clusters and their cognate mRNA targets, highlighting pseudogene insertions as a source of piRNA-guided gene silencing.

Our systematic analyses revealed that pachytene piRNAs regulate meiotic gene expression by establishing one-to-one relationships between individual piRNA clusters and their corresponding target genes. mRNA-targeting piRNAs arise from antisense pseudogene fragments embedded within piRNA clusters, echoing the ancestral use of transposon-derived sequences in pre-pachytene clusters to enforce transposon restriction in pre-meiotic germ cells ^4,19,22,23^. mRNA-targeting pachytene piRNAs form extensive swarms that engage their cognate mRNAs with near-perfect complementarity across broad regions, reminiscent of siRNA-mediated silencing yet distinct from the seed-based recognition typical of individual miRNAs^25^. Although siRNAs differ in biogenesis and act through AGO rather than PIWI proteins, both pathways employ slicer-dependent RNA degradation for post-transcriptional silencing, underscoring conserved mechanistic principles across distinct small RNA-silencing pathways.

To directly test the regulatory function of individual gene-targeting piRNA clusters, we engineered precise deletions of *piC-as(Spin1)* and *piC-as(Ago2)* in mice using CRISPR-Cas9 genome editing. Deletion of 668 bp spanning the *piC-as(Ago2)* promoter produced a loss-of-function allele that abolished all piRNAs from this locus (**Fig. 2a**). Despite this, *piC-as(Ago2)* knockout mice were fertile and displayed all stages of spermatogenesis, from spermatogonia through meiotic spermatocytes to spermatids and mature spermatozoa (**Fig. 2b, Extended Data Fig. 2a-c**). Notably, *Ago2* mRNA was the only transcript significantly upregulated in *piC-as(Ago2)* knockout testes (**Fig. 2c**). Transposon expression, piRNA precursor levels, and piRNA production from other clusters remained unchanged, indicating the absence of additional targets or cross-talk between *piC-as(Ago2)* and other clusters (**Fig. 2c, d**)^16^. The piRNAs from *piC-as(Ago2)* showed a strong preference for uridine at the first position (1U bias) -a hallmark of primary piRNA biogenesis^9,10^-but no enrichment for adenosine at position 10, suggesting that these piRNAs do not engage in ping-pong amplification (**Fig. 2e and Extended Data Fig. 2d**)^26^.

**Figure 2.**
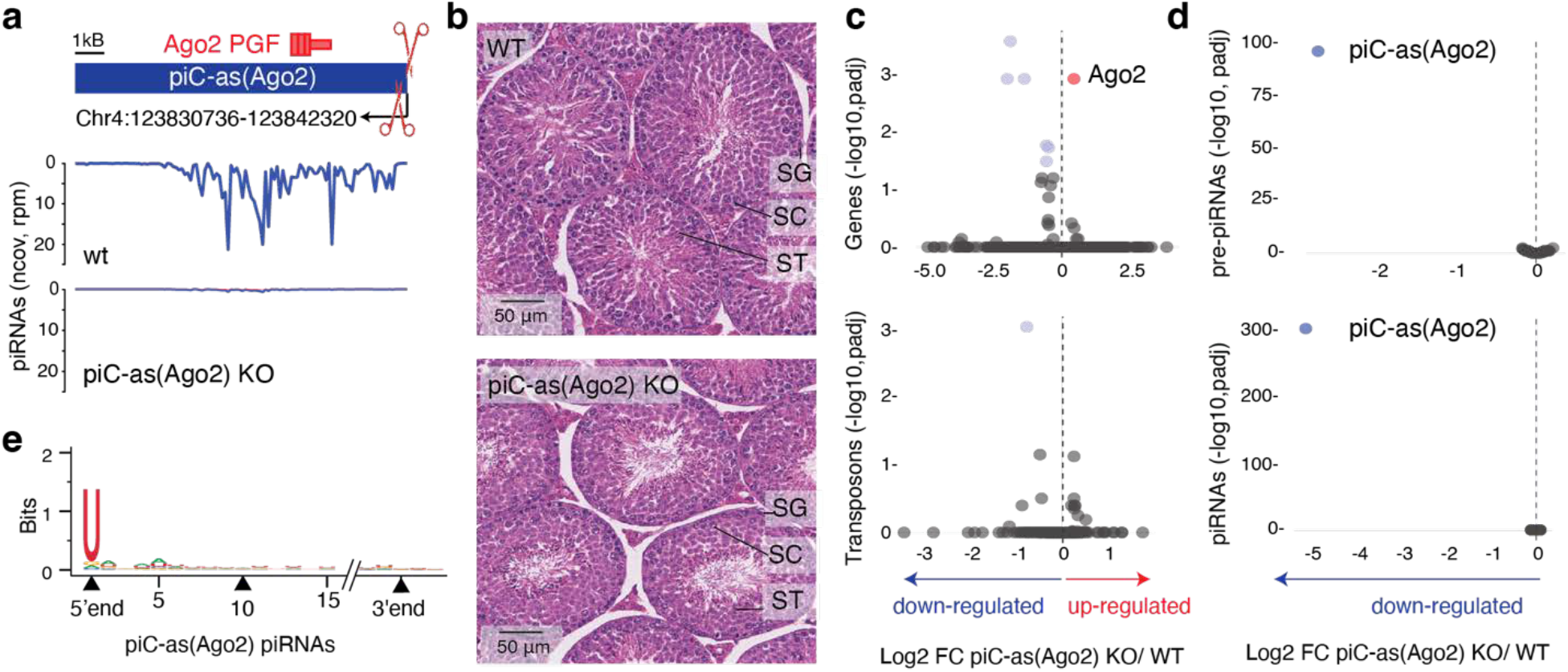
Loss of *piC-as(Ago2)* derepresses Ago2 during spermatogenesis. **a**, Schematic of *piC-as(Ago2)*, which generates *Ago2*-targeting piRNAs from the minus strand. A 668-bp promoter-spanning deletion was introduced by CRISPR–Cas9/sgRNAs to create a knockout (KO) allele. The *Ago2* pseudogene fragment (PGF) resides within the cluster (top, plus strand). Small-RNA-seq coverage (100-nt tiles; reads per million, rpm) shows loss of piRNAs in KO testes (bottom). **b**, Haematoxylin and eosin (H&E)–stained testis sections from wild-type (WT) and KO males show normal progression of spermatogenesis in both genotypes. SG, spermatogonia; SC, spermatocytes; ST, spermatids. **c**, Volcano plots of whole-testis RNA-seq showing log_2_ fold change (FC) in expression for protein-coding genes (>200 nt; RefSeq, top) and transposable elements (>200 nt; UCSC, bottom) in KO versus WT. *Ago2* is the only significantly upregulated gene. **d**, Volcano plots for RNA-seq (top) and small-RNA-seq (bottom) showing log_2_ FC for the top 64 meiotic piRNA clusters (90th percentile). piC-as(*Ago2*) is the only cluster with altered expression. RNA-seq: n = 3 biological replicates per genotype; small-RNA-seq: n = 2 per genotype. Significantly changed features (adjusted *P* < 0.05, two-sided; Benjamini–Hochberg) are coloured red (up) and blue (down). Dashed line indicates log_2_ FC = 0. **e**, Sequence logo of piC-as(*Ago2*) piRNAs shows a 1U bias and no enrichment for adenosine at position 10 (10A), consistent with primary piRNA biogenesis without ping-pong amplification.

Our results demonstrate that *piC-as(Ago2)* regulates a single dominant target gene, likely explaining the previously reported *Ago2* upregulation in *Miwi*-deficient spermatocytes^22^. This finding establishes a mechanistic paradigm for pachytene piRNA-guided gene regulation, in which individual piRNA clusters act as discrete regulatory units that control single major mRNA targets through an RNAi-like effector mechanism. The absence of overt spermatogenic defects in *piC-as(Ago2)* knockout mice eliminates confounding secondary effects, providing an unambiguous view of this direct regulatory relationship. This outcome parallels previous loss-of-function phenotypes of other clusters, such as *pi6* and *pi17*^*16*^, leaving unresolved which specific piRNAs underlie the spermatogenic arrest observed in *Miwi* knockout animals.

To further define the physiological role of individual gene-targeting piRNA clusters, we examined *piC-as(Spin1)*. Deleting 646 base pairs encompassing its promoter region abolished piRNAs from this locus, generating a loss-of-function allele (**Extended Data Fig. 3a**). Despite normal testis weight, litter size, and overall fertility, *piC-as(Spin1)KO* testes displayed striking multifocal spermatogenic defects (**Fig. 3a, Extended Data Fig. 3b**,**c**). A second loss-of-function allele, in which only the *Spin1* pseudogene fragment was removed from the cluster, produced comparable defects, establishing *Spin1* as the principal functional target of *piC-as(Spin1)* piRNAs (**Fig. 3b**).

**Figure 3.**
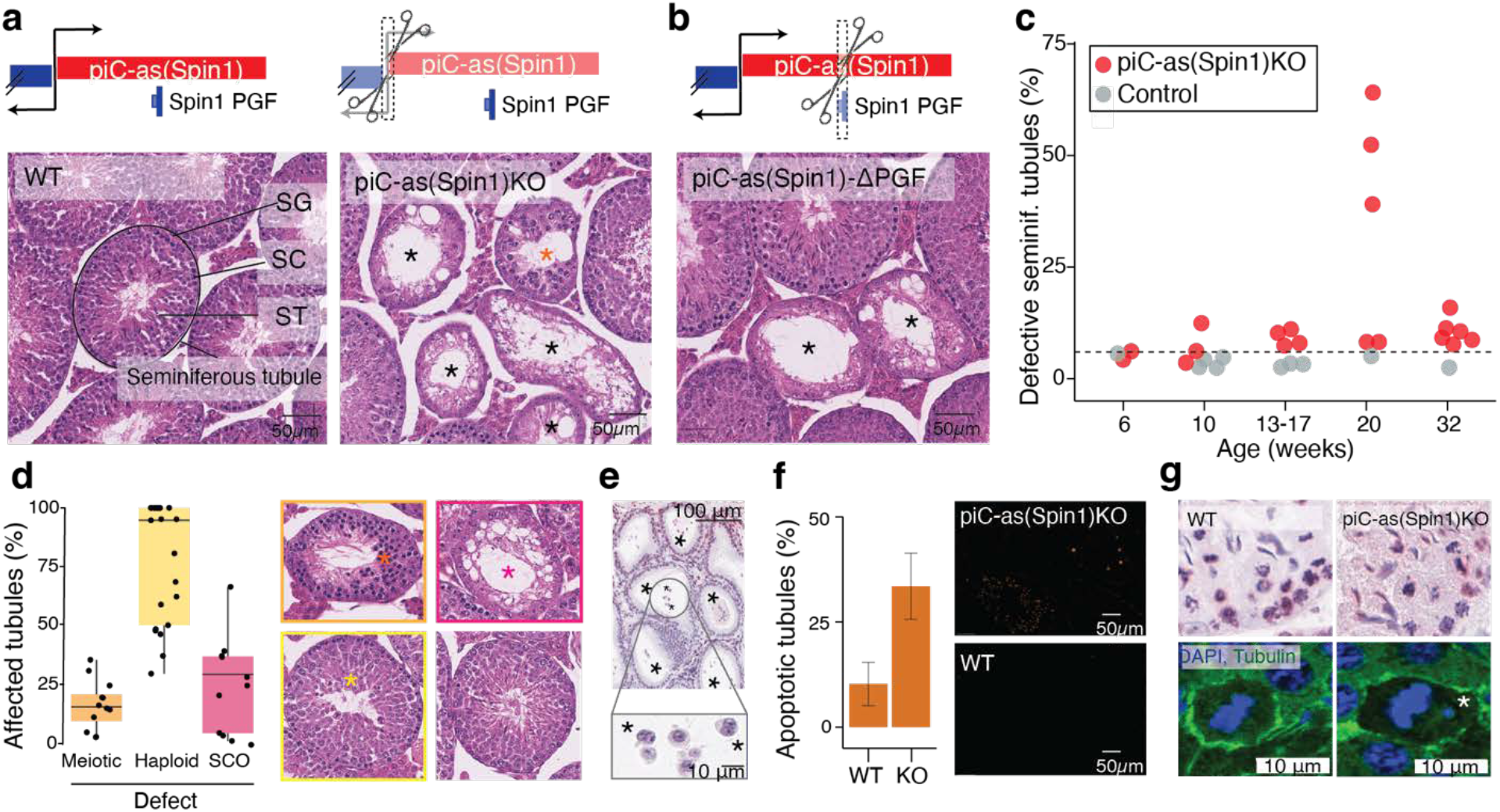
Loss of Spin1-targeting piRNAs causes metaphase misalignment and apoptosis during spermatogenesis. **a, b**, Schematic of the bidirectional piC-as(*Spin1*) locus. **a**, sgRNAs used to excise the promoter region to generate a KO allele. **b**, sgRNAs used to delete the *Spin1* pseudogene fragment (PGF). **Bottom (a**,**b):** representative haematoxylin and eosin (H&E)–stained testis sections. Both *piC-as(Spin1)* and *piC-as(Spin1)*-ΔPGF mutants show multifocal spermatogenic defects, including Sertoli cell-only (SCO) tubules (black asterisks); wild-type (WT) littermates appear normal. Orange asterisk indicates meiotic defect. SG, spermatogonia; SC, spermatocytes; ST, spermatids. **c**, Quantification of defective tubules per animal across age groups. Most *piC-as(Spin1)* KOs show pronounced defects compared with WT and heterozygous controls. Dashed line indicates the baseline frequency of naturally occurring defective tubules in healthy testes (6%). **d**, Composition of tubule defects per KO animal: fractions of Sertoli cell-only (SCO), meiotic and post-meiotic (haploid) abnormalities (colour-coded as in H&E images). Each animal contributes one or more points depending on defect types. In total, 5,333 tubules were scored from 21 KO animals. **e**, Representative H&E-stained epididymis from a severely affected KO showing absence of mature sperm and defective multinucleated spermatids (asterisks). **f**, TUNEL assay (TdT end-labelling with digoxigenin) shows increased apoptosis in KO versus WT tubules. Two WT and three KO animals were analysed; median proportions of TUNEL-positive tubules were 10% (WT) and 34% (KO). **g**, Representative H&E-stained seminiferous tubules showing misaligned chromosomes at metaphase in KO compared with proper alignment in WT (top). Immunofluorescence (DAPI, DNA; α-tubulin, spindle) confirms chromosome mispositioning at the spindle tip in KO tubules (bottom).

Systematic analysis across mice of different ages revealed variable penetrance of the spermatogenic defects, with <10% to >65% of seminiferous tubules affected in adult males (older than 13 weeks) (**Fig. 3c**). To rule out an impact of genetic modifiers, genome-wide monitoring using 10,000 marker genes confirmed that the extensively backcrossed line maintained a pure C57BL/6J background (**see Methods**). Notably, histopathological severity varied not only across individuals but even between testes of the same animal: one testis could appear normal while the contralateral testis exhibited extensive germ-cell loss and complete absence of mature sperm in the epididymis, indicating unilateral sterility (**Extended Data Fig. 3d**).

Histological examination revealed a spectrum of defects ranging from meiotic and post-meiotic (haploid) arrest to seminiferous tubules devoid of any germ cells, culminating in a severe Sertoli-cell-only (SCO) phenotype (**Fig. 3d**). The most prominent abnormalities occurred in post-meiotic spermatids, which frequently failed to elongate or complete cytokinesis, forming multinucleated cells. These defective post-meiotic cells were positive for γH2AX, indicative of chromatin damage (**Extended Data Fig. 3e**) and accumulated within the epididymis, where mature sperm are normally stored (**Fig. 3e**).

TUNEL assays revealed elevated apoptosis in 34% of knock-out seminiferous tubules compared with 10% in wild-type testes (**Fig. 3f**). Gene expression analysis excluded a contribution of unleashed transposons to the variable apoptotic phenotype (**Extended Data Fig. 3f**). In addition, *piC-as(Spin1) KO* testes exhibited chromosome mis-segregation defects reminiscent of those previously observed in *Miwi* and *Btbd18* knockout mice, both of which disrupt the pachytene piRNA pathway at large, including *piC-as(Spin1)* function (**Fig. 3g**)^3,27^. These findings indicate a phenotypic contribution of *piC-as(Spin1)* piRNAs reflected in broader piRNA-pathway mutants.

In summary, deletion of a single pachytene piRNA cluster targeting *Spin1* produced a multifocal spermatogenic phenotype marked by meiotic and post-meiotic defects and increased apoptosis, likely reflecting cumulative cellular damage that ultimately manifests as Sertoli-cell-only tubules. The observed apoptosis and chromosome mis-segregation mirror *Spin1* overexpression phenotypes reported in oocytes and somatic cells^28,29^. Together, these findings challenge the prevailing view that individual piRNA clusters are dispensable for spermatogenesis^16,30^ and suggest that the *Miwi* knockout phenotype may reflect the combined outcome of multiple, partially penetrant defects arising from the loss of distinct gene-regulatory piRNA clusters.

Next, we asked whether piRNA-guided meiotic gene regulation is conserved in humans. Similar to our observations in mouse (**Fig. 1**), we found that ∼2.7% of human pachytene piRNAs target mRNAs in trans with near-perfect complementarity across extended regions of coding sequence and 3’UTR (**Fig. 4a,b, Extended Data Fig. 4a**). The top target, *GOLGA2* accumulated nearly 15% of all mRNA-targeting piRNAs across more than 2 kilobases (**Fig. 4b, Extended Data Fig. 4b**), and most its targeting piRNAs derived from a single cluster, *piC-as(GOLGA2*), containing a GOLGA2 pseudogene fragment (**Fig. 4c, Extended Data Fig. 4c**,**d**). These findings establish that the phenomenon of meiotic piRNA-guided gene targeting is conserved, though the specific mRNA targets differ between mouse and human. The lack of target conservation is consistent with the rapid evolution of piRNAs, piRNA clusters, and their pre-piRNA transcripts, which have been reported to be among the fastest-evolving genomic sequences^5^, and most likely reflect quickly evolving germ cell biology.

**Figure 4.**
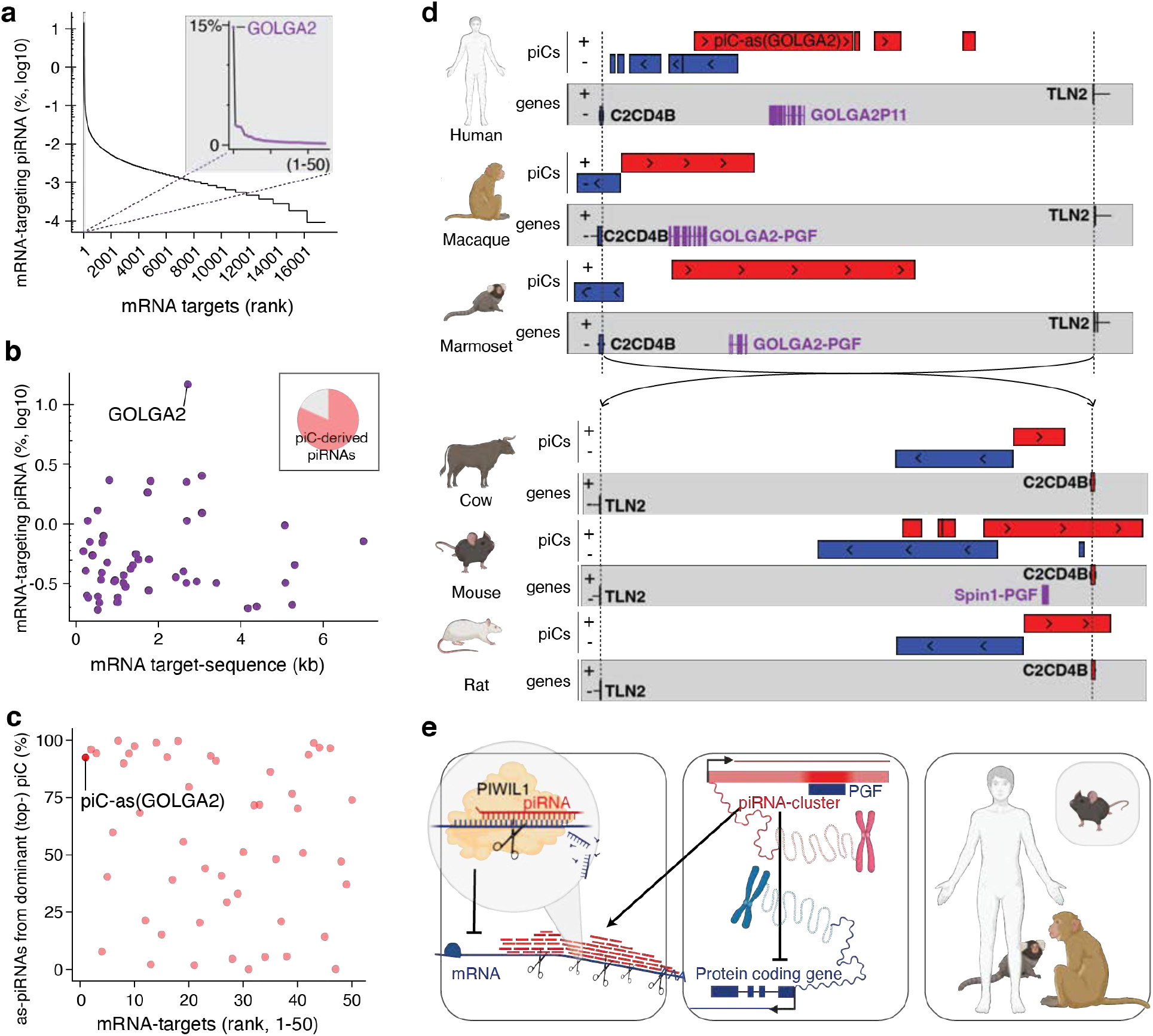
Gene-targeting pachytene piRNAs define a conserved, RNAi-like mechanism of meiotic gene control in mammals. **a**, Human mRNA targets ranked by pachytene-piRNA coverage. **Inset:** top 50 targets; the top-ranked gene, *GOLGA2*, is highlighted (purple). **b**, Relationship between mRNA length (kb) and the fraction of targeting piRNAs (log_10_ of % among all human mRNA-targeting piRNAs) for the top 50 targets; *GOLGA2* is indicated. **Inset:** proportion of trans-acting piRNAs (targets outside their cluster locus) originating from piRNA clusters (81.7%); *GOLGA2* targeting is dominated by piRNAs from *piC-as(GOLGA2)*. **c**, For each gene, fraction of targeting piRNAs contributed by the dominant (top) piRNA cluster; *piC-as(GOLGA2)* is labelled. **d**, Synteny maps of pachytene piRNA clusters (piCs) and neighbouring genes (*C2CD4B* and *TLN2*) across mammals. In primates (human, macaque, marmoset), *GOLGA2*-derived pseudogene fragments (PGFs; purple) lie antisense within a syntenic piC region, suggesting a conserved regulatory role. In rodents, syntenic piCs occur in mouse and rat; only mouse carries the *Spin1*-derived PGF corresponding to *piC-as(Spin1)*. In cow, the syntenic piC is retained despite the absence of the PGF, suggesting that the piC predates PGF acquisition in a common ancestor. Regions are aligned by *C2CD4B*/*TLN2* boundaries (dashed line). Red, plus strand; blue, minus strand. **e, Model:** Groups of pachytene piRNAs collectively target mRNAs across extended regions evoking the specificity and efficacy of siRNA-guided gene silencing. (left). Pseudogene fragments (PGF) within piRNA clusters (piCs) generate piRNAs with near-perfect complementarity to their cognate mRNAs. They establish a one-to-one correspondence between a piRNA cluster and a single target gene (middle). This cluster–gene control principle is conserved across mammals, supporting a unifying model in which piRNA-guided silencing has been co-opted for meiotic gene regulation. Whereas most cluster–gene pairs evolve rapidly, a primate-specific piRNA cluster targeting *GOLGA2* has been conserved for more than 10 million years (see also Extended Data Figure 4e). (human, rhesus macaque, marmoset, and mouse; right)

Although piRNA sequences are generally not conserved, some meiotic piRNA clusters are syntenic across mammals^5^. To test whether *piC-as(GOLGA2)* has syntenic counterparts in non-human primates, we analyzed rhesus macaque and marmoset, both of which have experimentally defined piRNA clusters^5,31^. Remarkably, both species contained a *GOLGA2* fragment within the syntenic cluster suggesting potential conservation of its regulatory function (**Fig. 4d**). Moreover, *piC-as(GOLGA2)* shared more than 80% sequence identity between human and macaque, challenging the prevailing paradigm that piRNA clusters lack sequence conservation across different species (**Extended Data Fig. 4e**).

Extending our analysis across mammals, we identified a syntenic piRNA cluster in cow, rat and mouse (**Fig. 4d**). These loci lack the GOLGA2-derived pseudogene fragment, indicating that the cluster predated pseudogene insertion. A striking link emerged in mouse: the syntenic cluster corresponds to piC-as(*Spin1*), whose regulatory role in spermatogenesis we established above (**Fig. 4d; Extended Data Fig. 4e**). Phylogenetic mapping places the origin of this syntenic cluster >90 million years ago and supports two independent co-option events mediated by pseudogene insertions, one producing the *Spin1*-targeting regulon in mice, and another generating a conserved *GOLGA2*-targeting regulon across primates.

Acquisition of regulatory pseudogene fragments by an ancestral piRNA cluster suggests a mechanism distinct from the de novo formation of protective pre-pachytene clusters during retroviral invasion. In the latter, new piRNA clusters arise via readthrough transcription to capture viral sequence information downstream of genes (piC-DoGs)^23,32^. Instead, varying pseudogene fragments in pachytene piRNA clusters suggests the evolutionary repurposing of existing piRNA clusters, echoing co-option phenomena best described for ancient retroviruses and other transposons that gave rise to novel protein-coding genes and regulatory elements^33^. These data propose a co-option model in which a duplication of an ancestral genome-defence pathway was repurposed for meiotic gene regulation.

Overall, our results define a molecular framework for pachytene piRNA function, resolve the underlying mechanism, and advance an evolutionary model (**Fig. 4e**). By shifting our focus from single piRNAs to groups of piRNAs, we reveal an RNAi-like mode of gene regulation in which thousands of piRNAs from one cluster converge on a single mRNA, forming a one-to-one cluster–gene regulon. Targeted deletion of a single piRNA cluster or its resident pseudogene fragment eliminates the cognate piRNAs, derepresses the major target mRNA, and causes spermatogenic defects in mice. Comparative genomics delineates a conserved meiotic programme in mammals alongside rapidly evolving cluster–target pairs, and identifies a uniquely conserved primate regulon, piC-as(GOLGA2). Synteny between this cluster and piC-as(Spin1) in mouse points to a mechanism of piRNA cluster formation distinct from pre-meiotic clusters that arise downstream of genes (piC-DoGs). Together, these findings recast pachytene piRNAs as gene-specific regulators of meiosis and lay the foundation for defining when, where, and under what selective pressures this programme was installed during mammalian evolution.

## Acknowledgements

We thank members of the Haase, Macfarlan, Meister, Hafner, Pratto, and Dean laboratories, for helpful discussions. We are grateful to NIH core facilities and animal programs for expert assistance, in particular the NIDDK Animal Program (Michael Eichner, Oksana Gavrilova and Jennifer Portas), the NHLBI Light Microscopy Core (Christian Combs), the NIAMS Genomics Core (Stefania Dell’Orso and Faiza Naz), and the NIH HPC (Biowulf). F.A. is supported by an Intramural AIDS Research Fellowship (AIRF). This work was supported by the Intramural Research Program of the NIH, NIDDK (ZIA DK075111 to A.D.H.).

## Ethics declaration

The authors declare no competing interests.

## Data availability

All data supporting the findings are available within the paper and its Supplementary Information or from public repositories as indicated. Newly generated mouse lines will be available from The Jackson Laboratory under assigned JAX stock numbers and RRIDs upon publication. RNA and piRNA sequencing data are available through the European Nucleotide Archive (ENA) PRJEB102223 (ERP183623).

## Code availability

Custom code used in this study is provided as Supplementary Code and will also be released on GitHub and archived on Zenodo upon publication.

## Author contributions

Z.L., F.A., D.S. and A.D.H. designed the study, performed experiments, analysed data and wrote the manuscript. D.S. conceived the target-centred analysis. Z.L. led all mouse work and phenotypic characterisation, performed sequencing, and generated and analysed the data for Figs. 2–3. F.A. performed most analyses for Figs. 1 and 4. R.C. and T.M. carried out the evolutionary-distance analyses (Fig. 4). S.R. and T.M. provided technical support for generation of the initial mouse line. G.M. co-mentored F.A. All authors discussed the results and contributed to the final manuscript.

Supplementary Information is available for this manuscript.

Correspondence and requests for materials should be addressed to Z.L. and A.D.H..

## References

1. Czech B, Munafo M, Ciabrelli F, et al. piRNA-Guided Genome Defense: From Biogenesis to Silencing. Annu Rev Genet 2018;52:131–157. DOI: 10.1146/annurev-genet-120417-031441.

2. Stallmeyer B, Buhlmann C, Stakaitis R, et al. Inherited defects of piRNA biogenesis cause transposon de-repression, impaired spermatogenesis, and human male infertility. Nat Commun 2024;15(1):6637. DOI: 10.1038/s41467-024-50930-9.

3. Deng W, Lin H. miwi, a murine homolog of piwi, encodes a cytoplasmic protein essential for spermatogenesis. Dev Cell 2002;2(6):819–30. DOI: 10.1016/s1534-5807(02)00165-x.

4. Ozata DM, Gainetdinov I, Zoch A, O’Carroll D, Zamore PD. PIWI-interacting RNAs: small RNAs with big functions. Nat Rev Genet 2019;20(2):89–108. DOI: 10.1038/s41576-018-0073-3.

5. Ozata DM, Yu T, Mou H, et al. Evolutionarily conserved pachytene piRNA loci are highly divergent among modern humans. Nat Ecol Evol 2020;4(1):156–168. DOI: 10.1038/s41559-019-1065-1.

6. Joshua-Tor L, Hannon GJ. Ancestral roles of small RNAs: an Ago-centric perspective. Cold Spring Harb Perspect Biol 2011;3(10):a003772. DOI: 10.1101/cshperspect.a003772.

7. Fu Q, Wang PJ. Mammalian piRNAs: Biogenesis, function, and mysteries. Spermatogenesis 2014;4:e27889. DOI: 10.4161/spmg.27889.

8. Aravin AA, Hannon GJ, Brennecke J. The Piwi-piRNA pathway provides an adaptive defense in the transposon arms race. Science 2007;318(5851):761–4. DOI: 10.1126/science.1146484.

9. Aravin A, Gaidatzis D, Pfeffer S, et al. A novel class of small RNAs bind to MILI protein in mouse testes. Nature 2006;442(7099):203–7. DOI: 10.1038/nature04916.

10. Girard A, Sachidanandam R, Hannon GJ, Carmell MA. A germline-specific class of small RNAs binds mammalian Piwi proteins. Nature 2006;442(7099):199–202. DOI: 10.1038/nature04917.

11. Watanabe T, Chuma S, Yamamoto Y, et al. MITOPLD is a mitochondrial protein essential for nuage formation and piRNA biogenesis in the mouse germline. Dev Cell 2011;20(3):364–75. DOI: 10.1016/j.devcel.2011.01.005.

12. Ipsaro JJ, Haase AD, Knott SR, Joshua-Tor L, Hannon GJ. The structural biochemistry of Zucchini implicates it as a nuclease in piRNA biogenesis. Nature 2012;491(7423):279–83. DOI: 10.1038/nature11502.

13. Reuter M, Berninger P, Chuma S, et al. Miwi catalysis is required for piRNA amplification-independent LINE1 transposon silencing. Nature 2011;480(7376):264–7. DOI: 10.1038/nature10672.

14. Vourekas A, Zheng Q, Alexiou P, et al. Mili and Miwi target RNA repertoire reveals piRNA biogenesis and function of Miwi in spermiogenesis. Nat Struct Mol Biol 2012;19(8):773–81. DOI: 10.1038/nsmb.2347.

15. Gainetdinov I, Vega-Badillo J, Cecchini K, et al. Relaxed targeting rules help PIWI proteins silence transposons. Nature 2023;619(7969):394–402. DOI: 10.1038/s41586-023-06257-4.

16. Wu PH, Fu Y, Cecchini K, et al. The evolutionarily conserved piRNA-producing locus pi6 is required for male mouse fertility. Nat Genet 2020;52(7):728–739. DOI: 10.1038/s41588-020-0657-7.

17. Genzor P, Konstantinidou P, Stoyko D, et al. Cellular abundance shapes function in piRNA-guided genome defense. Genome Res 2021;31(11):2058–2068. DOI: 10.1101/gr.275478.121.

18. Gainetdinov I, Colpan C, Arif A, Cecchini K, Zamore PD. A Single Mechanism of Biogenesis, Initiated and Directed by PIWI Proteins, Explains piRNA Production in Most Animals. Mol Cell 2018;71(5):775–790 e5. DOI: 10.1016/j.molcel.2018.08.007.

19. Dias Mirandela M, Zoch A, Leismann J, et al. Two-factor authentication underpins the precision of the piRNA pathway. Nature 2024;634(8035):979–985. DOI: 10.1038/s41586-024-07963-3.

20. Anzelon TA, Chowdhury S, Hughes SM, Xiao Y, Lander GC, MacRae IJ. Structural basis for piRNA targeting. Nature 2021;597(7875):285–289. DOI: 10.1038/s41586-021-03856-x.

21. Dowling M, Homolka D, Raad N, Gos P, Pandey RR, Pillai RS. In vivo PIWI slicing in mouse testes deviates from rules established in vitro. RNA 2023;29(3):308–316. DOI: 10.1261/rna.079349.122.

22. Watanabe T, Cheng EC, Zhong M, Lin H. Retrotransposons and pseudogenes regulate mRNAs and lncRNAs via the piRNA pathway in the germline. Genome Res 2015;25(3):368–80. DOI: 10.1101/gr.180802.114.

23. Konstantinidou P, Loubalova Z, Ahrend F, et al. A comparative roadmap of PIWI-interacting RNAs across seven species reveals insights into de novo piRNA-precursor formation in mammals. Cell Rep 2024;43(10):114777. DOI: 10.1016/j.celrep.2024.114777.

24. Kazazian HH, Jr., Moran JV. Mobile DNA in Health and Disease. N Engl J Med 2017;377(4):361–370. DOI: 10.1056/NEJMra1510092.

25. Bartel DP. Metazoan MicroRNAs. Cell 2018;173(1):20–51. DOI: 10.1016/j.cell.2018.03.006.

26. Aravin AA, Sachidanandam R, Girard A, Fejes-Toth K, Hannon GJ. Developmentally regulated piRNA clusters implicate MILI in transposon control. Science 2007;316(5825):744–7. DOI: 10.1126/science.1142612.

27. Zhou L, Canagarajah B, Zhao Y, et al. BTBD18 Regulates a Subset of piRNA-Generating Loci through Transcription Elongation in Mice. Dev Cell 2017;40(5):453–466 e5. DOI: 10.1016/j.devcel.2017.02.007.

28. Yuan H, Zhang P, Qin L, et al. Overexpression of SPINDLIN1 induces cellular senescence, multinucleation and apoptosis. Gene 2008;410(1):67–74. DOI: 10.1016/j.gene.2007.11.019.

29. Choi JW, Zhao MH, Liang S, et al. Spindlin 1 is essential for metaphase II stage maintenance and chromosomal stability in porcine oocytes. Mol Hum Reprod 2017;23(3):166–176. DOI: 10.1093/molehr/gax005.

30. Choi H, Wang Z, Dean J. Sperm acrosome overgrowth and infertility in mice lacking chromosome 18 pachytene piRNA. PLoS Genet 2021;17(4):e1009485. DOI: 10.1371/journal.pgen.1009485.

31. Hirano T, Iwasaki YW, Lin ZY, et al. Small RNA profiling and characterization of piRNA clusters in the adult testes of the common marmoset, a model primate. RNA 2014;20(8):1223–37. DOI: 10.1261/rna.045310.114.

32. Yu T, Blyton MBJ, Abajorga M, et al. Evolution of KoRV-A transcriptional silencing in wild koalas. Cell 2025;188(8):2081–2093 e16. DOI: 10.1016/j.cell.2025.02.006.

33. Chuong EB, Elde NC, Feschotte C. Regulatory activities of transposable elements: from conflicts to benefits. Nat Rev Genet 2017;18(2):71–86. DOI: 10.1038/nrg.2016.139.

