## Supplemental Figures for "Pachytene piRNAs define a conserved program of meiotic gene regulation": 1_Loubalova_2025_bX_SUP.pdf

### Extended Data Figures

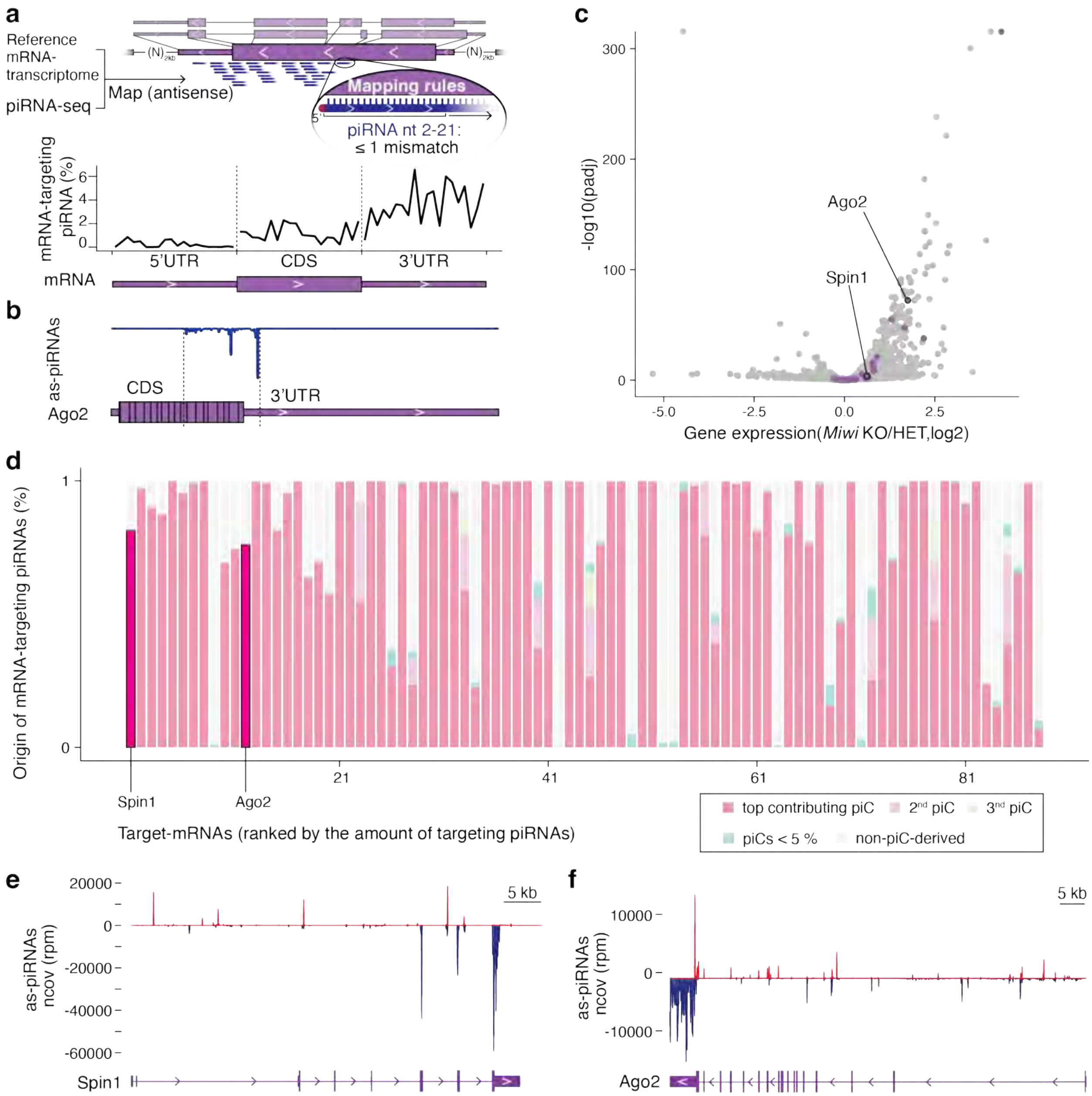


**Extended Data Fig. 1 | Gene-targeting pachytene piRNAs establish a one-to-one regulatory relationship between piRNA clusters and cognate target mRNAs. a,** Target-centered search for piRNA–mRNA pairs (≤1 mismatch across nt 2–21) using antisense mapping to a custom mRNA transcriptome identifies mRNAs hit by many piRNAs across broad sequence intervals. This approach recovers mismatch-mediated targeting that genome-based mapping misses (which assigns piRNAs uniquely to their source clusters). Metagene plot shows the distribution of piRNA 5′ ends across 5′ UTR, CDS and 3′ UTR (each feature scaled to 20 bins; averaged over the 88 top-targeted mRNAs from Fig. 1a inset), revealing a bias toward the 3′ UTR. **b,**

Antisense piRNA coverage across *Ago2* exons (piRNA density, blue) overlaid on the *Ago2* gene model (purple). **c,** Volcano plot of differential expression in *Miwi* knockout (KO) versus heterozygous (HET) spermatocytes; x-axis, $\log_2$ fold change; y-axis, $-\log_{10}$(adjusted *P*). Top-targeted genes, including *Spin1* and *Ago2*, are highlighted (purple). **d,** Contribution of pachytene piRNA clusters (piCs) to the pool of targeting piRNAs for the top-ranked genes. Each bar represents one gene; colours denote the dominant (top), second and third contributing piCs, minor piCs (<5% each) and non-piC loci. *Spin1* and *Ago2* are labelled. **e,** Antisense piRNA coverage across *Spin1* (piRNA density: red, plus strand; blue, minus strand) over the *Spin1* gene model (purple), normalized to reads per million (rpm) within the displayed region. **f,** As in **e** for *Ago2*.

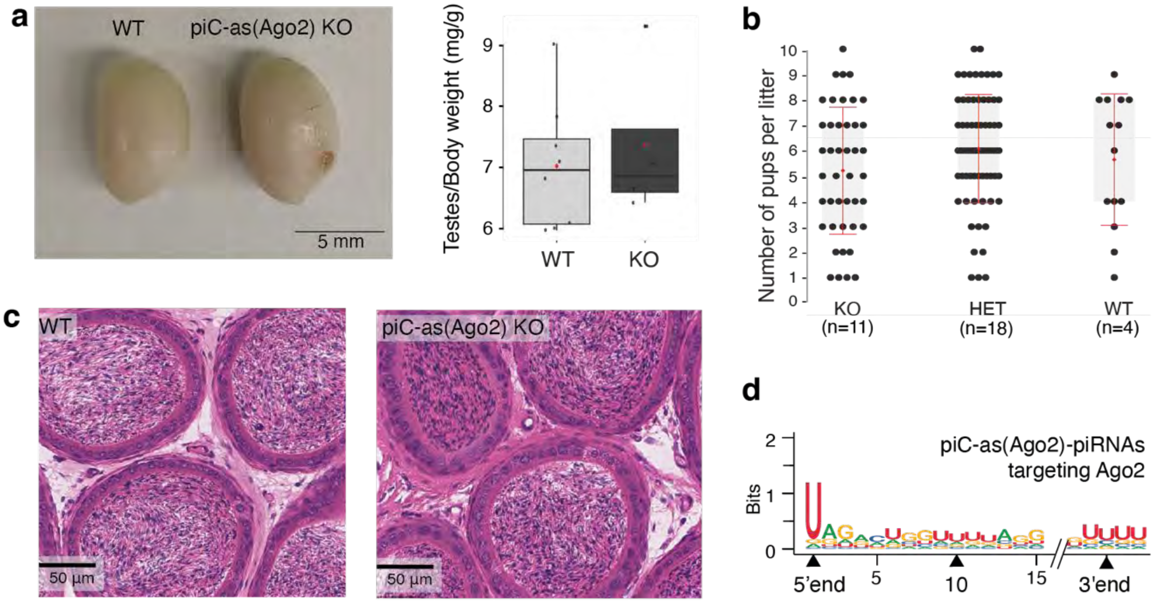


**Extended Data Fig. 2 | Loss of *piC-as(Ago2)* derepresses Ago2 without impairing spermatogenesis.** **a,** Representative testes (left) and testis:body-weight ratios (right) for wild-type (WT) and *piC-as(Ago2)* knockout (KO) males show no significant difference. **b,** Breeding performance: average litter sizes are comparable among KO, heterozygous and WT animals. **c,** Haematoxylin and eosin (H&E)–stained epididymides from WT and KO males show abundant mature sperm in both genotypes. **d,** Sequence logo of *piC-as(Ago2)*–derived piRNAs shows a strong 1U bias and no enrichment at position 10, consistent with primary piRNA biogenesis without ping-pong amplification.

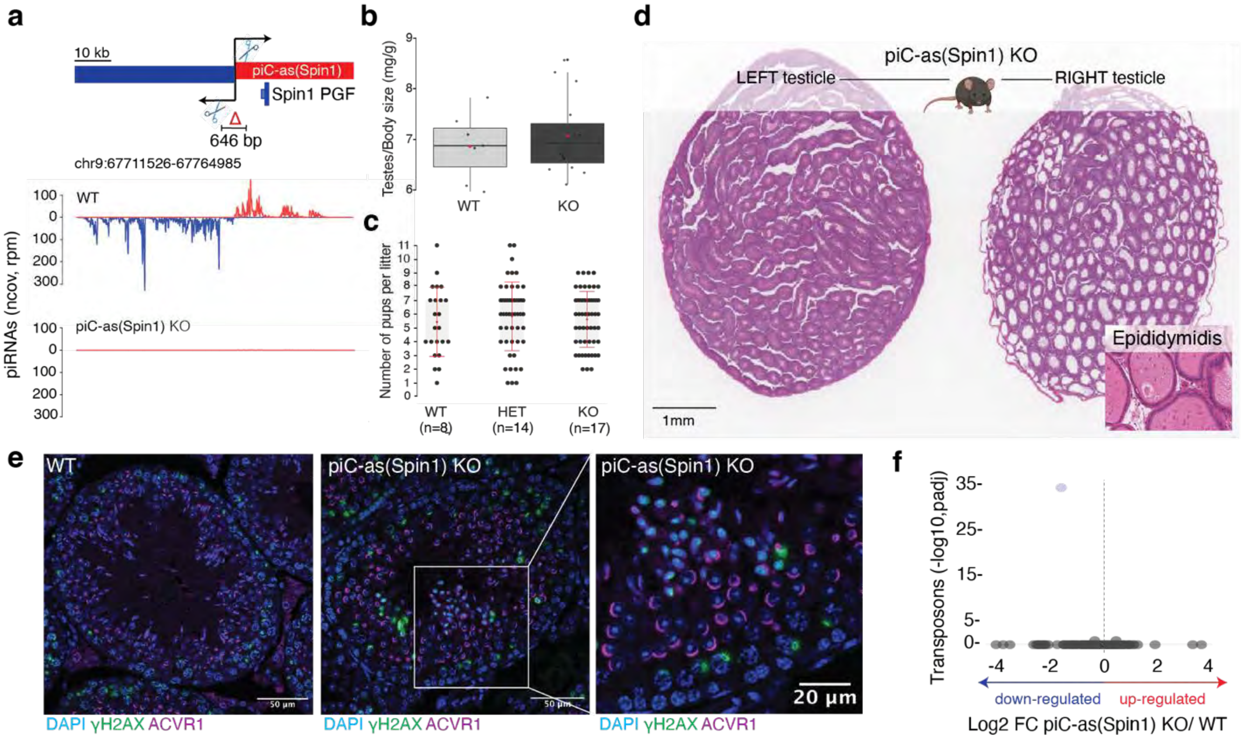


**Extended Data Fig. 3 | Loss of *Spin1*-targeting piRNAs causes multifocal spermatogenic defects with variable penetrance.** **a,** Schematic of the bidirectional *piC-as(Spin1)* locus showing the CRISPR–Cas9 excision within the predicted promoter and the position of the antisense *Spin1* pseudogene fragment (PGF) in the cluster (top). Small-RNA-seq coverage (100-nt tiles; reads per million, rpm) shows loss of piRNAs in *piC-as(Spin1)* knockout (KO) testes (bottom). **b,** Testis:body-weight ratios do not differ significantly between KO and wild-type (WT) males. **c,** Breeding performance: average litter sizes are comparable among KO, heterozygous and WT animals. **d,** Haematoxylin and eosin (H&E)–stained contralateral testes from a 20-week-old KO male showing a healthy left testis and a severely affected right testis with >60% of tubules displaying increased luminal diameter. **Inset:** H&E-stained epididymis attached to the affected testis shows degenerating cells in place of mature sperm, consistent with unilateral sterility. **e,** Immunofluorescence of seminiferous tubules from WT (left) and KO (middle, right): DAPI (DNA, blue), γH2AX (DNA double-strand break marker, green) and ACVR1 (post-meiotic spermatids, magenta). Magnified view (right) highlights γH2AX-positive post-meiotic cells, indicating compromised chromatin integrity in KO testes. **f,** Volcano plot of whole-testis RNA-seq showing $\log_2$ fold change for transposable elements (>200 nt; UCSC annotation) in KO versus WT; no TE upregulation is detected in *piC-as(Spin1)* KO testes.

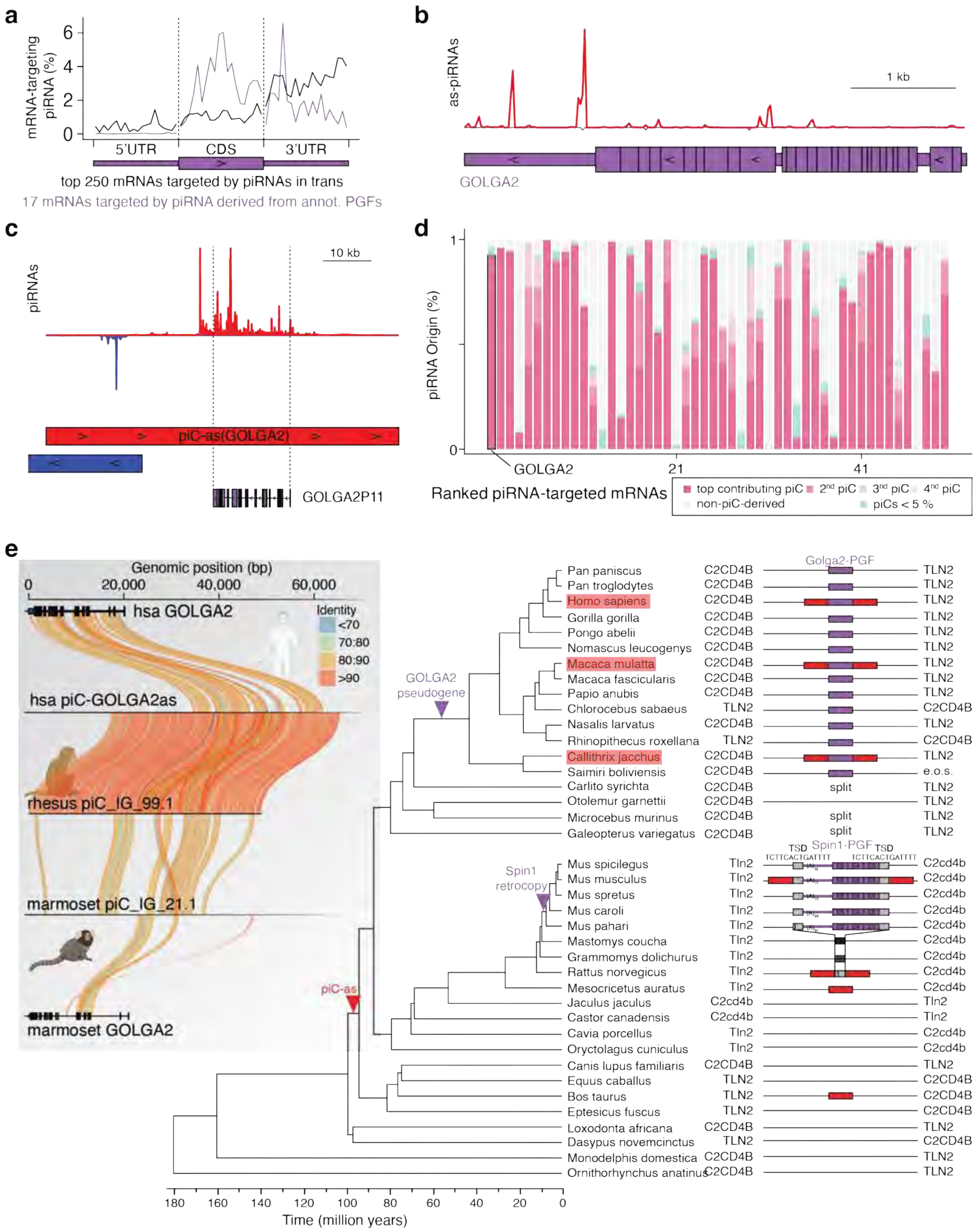


**Extended Data Fig. 4 | A conserved piRNA cluster targeting *GOLGA2* in primates and its pseudogene-guided architecture. a,** Distribution of mRNA-targeting piRNAs across transcript features (5′ UTR, CDS and 3′ UTR) for the top 250 targets in human (black) and for the subset (n = 17) whose targeting piRNA clusters (piCs) contain the cognate pseudogene fragment (PGF; purple). Each feature was scaled to 20 bins per gene; values are the mean fraction of piRNA 5′ ends per bin. **b,** Antisense piRNA coverage across collapsed *GOLGA2* exons (piRNA density, red) overlaid on the *GOLGA2* gene model (purple). **c,** piRNA coverage at *piC-as(GOLGA2)* highlighting the *GOLGA2* pseudogene (GOLGA2P11, purple) with antisense piRNAs (chr15:62,207,115–62,280,150; red, plus strand; blue, minus strand). **d,** Contribution of individual piCs to the pool of targeting piRNAs for the top targets. Each bar represents one gene; colours denote the dominant (top), second and third contributing piCs, minor piCs (<5%), and non-piC loci. *GOLGA2* is highlighted. **e,** Phylogeny of a piC located between *TLN2* and *C2CD4B* across mammals. Species with empirically detected piRNAs are marked (red boxes). The *Spin1* PGF (purple), flanked by 14-bp target-site duplications (TSDs), is present in all *Mus* species but absent in related rodents (empty site; single target site); some non-*Mus* rodents carry a mutated site (black bars). The *GOLGA2* PGF (purple) is present in most primates, including New World and Old World monkeys, lesser apes and great apes. Approximate timings of piC formation and PGF insertion were inferred by parsimony (e.o.s., end of scaffold; split, *C2CD4B* and *TLN2* on separate scaffolds). **Inset (grey):** pairwise identity across the *GOLGA2*-associated piRNA locus in primates, comparing human *GOLGA2*, human *piC-as(GOLGA2)*, rhesus pi_IG_99.1, marmoset pi_IG_21.1 (ref) and marmoset *GOLGA2*; ribbons are coloured by percent identity; gene models show exon–intron structures (thick, CDS; thin, UTRs).
